# Modeling cell infection via virus-producing cells rather than free infectious virus significantly improves fits of *in vitro* viral kinetic data

**DOI:** 10.1101/627968

**Authors:** Veronika Bernhauerová, Veronica V. Rezelj, Laura I. Levi, Marco Vignuzzi

## Abstract

Chikungunya and Zika viruses are arthropod-borne viruses that pose significant threat to public health. Experimental data show that during *in vitro* infection both viruses exhibit qualitatively distinct replication cycle kinetics. Chikungunya viral load rapidly accumulates within the first several hours post infection whereas Zika virus begins to increase at much later times. We sought to characterize these qualitatively distinct *in vitro* kinetics of chikungunya and Zika viruses by fitting a family of mathematical models to time course viral load datasets. We demonstrate that the standard viral kinetic model, which considers that new infections result only from free virus penetrating susceptible cells, does not fit experimental data as well as a model in which the number of virus-infected cells is the primary determinant of infection rate. We provide biologically meaningful quantifications of the main viral kinetic parameters and show that our results support cell-to-cell or localized transmission as a significant contributor to viral infection with chikungunya and Zika viruses.

**Importance:** Mathematical modeling has become a useful tool to tease out information about virus-host interactions and thus complements experimental work in characterizing and quantifying processes within viral replication cycle. Importantly, mathematical models can fill in incomplete data sets and identify key parameters of infection, provided the appropriate model is used. The *in vitro* time course dynamics of mosquito transmitted viruses, such as chikungunya and Zika, have not been studied by mathematical modeling and thus limits our knowledge about quantitative description of the individual determinants of viral replication cycle. This study employs dynamical modeling framework to show that the rate at which cells become virus-infected is proportional to the number or virus-infected cells rather than free extracellular virus in the milieu, a widely accepted assumption in models of viral infections. Using the refined mathematical model in combination with viral load data, we provide quantification of the main drivers of chikungunya and Zika *in vitro* kinetics. Together, our results bring quantitative understanding of the basic components of chikungunya and Zika virus dynamics.

## Introduction

Chikungunya (CHIKV) and Zika (ZIKV) viruses are arthropod-borne viruses (arbovirus) primarily transmitted through a bite of infected *Aedes* mosquitoes, and their continuous re-emergence pose an important public health threat. CHIKV was originally isolated in 1953 during an epidemic outbreak in Tanzania [1]. Outbreaks of CHIKV occurred in the western Indian Ocean in 2005-6 [2], India and Italy in 2007 along with several Southeast Asian countries, Pacific regions and the Americas [3]. Similarly, ZIKV was first discovered in 1947 in a Ugandan forest [4]. The first sporadic ZIKV outbreaks outside Africa were reported in the Asia-Pacific region in 2007 [5] and 2013 [6], followed by its rapid spread to the Western hemisphere in 2016 [7], which received public attention due to the association of ZIKV infection with newborn microcephaly and other neurological abnormalities [8–11]. Currently, no approved vaccine or therapeutic treatments exist to specifically target CHIKV or ZIKV infections. Disease prevention mostly relies on decreasing the number of transmission events through vector control strategies, presenting a significant challenge to limit the incidence of future epidemics, especially in developing countries.

Although CHIKV and ZIKV belong to distinct virus families (*Togaviridae* and *Flaviviridae*, respectively), virus particles share common characteristics, such as their positive single-stranded RNA genome and the presence of a lipid envelope derived from the host. Both viruses infect a wide spectrum of mosquito and mammalian cell lines, including Vero cells, mosquito cells Aag2 or C6/36, as well as various human cell lines, including Huh7 [12, 13]. The classical kinetics following infection of a non-lytic virus begins with an eclipse phase in which attachment, entry and the first round of replication and assembly occurs. This period is followed by an exponential increase in viral particles released to the extracellular milieu following virus egress. Finally, a plateau phase is reached when the maximum capacity of virus production by the cells is reached. Following the plateau phase, the number of infectious virus particles in the extracellular milieu generally begins to decline due to a loss in stability and infectivity of the virions in the environment. Importantly, the time for each of these phases of virus replication kinetics may vary between virus types, strains, and cell type, as the rate of different processes occurring in an infected cell (such as penetration, uncoating, replication, budding) may differ under different conditions.

Mathematical models of *in vitro* viral infections help elucidate the time scales of each of these phases and characteristics affecting virus-infected cells, such as the length of eclipse phase, or the mean lifespan of virus-releasing cells. Dynamical models provide accurate estimations of the rates that dictate the accumulation of virus in the free space outside of cells, such as viral genome production rates and loss of viral infectivity. The more precise quantification of such fundamental processes within virus-host interactions can better replace generic, experiment-specific, qualitative descriptions of virus replication (e.g., ‘attenuated growth’, ‘reduced fitness’ *in vitro*). These measures have been determined for a number of viruses, including HIV-1 and simian-human immunodeficiency virus (SHIV) [14–19], hepatitis C virus [20–24], poliovirus [25–27], influenza A virus and its variants [28–32], West Nile virus [33], and Ebola virus [34, 35]. The mathematical models proposed in these experimental studies rely on the assumption that infection of susceptible cells occurs via free infectious virus. In contrast, only a limited number of theoretical studies have considered infection of susceptible cells to be proportional to the total number of virus-producing cells, which is commonly referred to as the cell-to-cell transmission model. The latter type of modeling is important to consider, since cell-to-cell viral transmission has been observed to be an additional contributor to the infection for many enveloped viruses [36–38]. Indeed, some theoretical studies showed that cell-to-cell transmission of virus contributed approximately equally to the *in vitro* growth of equine infectious anaemia virus [39] and HIV-1 [40], explained multiplicity of (HIV-1)-infected splenocytes in humans [41], or permitted spread of HIV-1 virus despite antiretroviral therapy [42, 43]. Interestingly, while no direct evidence of cell-to-cell transmission of ZIKV exists to date, indirect evidence of cell-to-cell transmission of CHIKV was previously suggested to enable CHIKV resistance to antibody neutralization by bypassing the extracellular space [44, 45].

Modeling cell-to-cell transmission most commonly refers to modeling direct biological transfer of a virus. However, modeling cell-to-cell transmission may also be viewed as a proxy to model localized infections caused by low amounts of free infectious virus. This may especially be enhanced in static conditions, such as cell culture, where cell infections are more likely to occur in a localized manner as viral particles produced by an infected cell penetrate susceptible cells within their immediate neighborhood. In addition, such relatively low amounts of infectious virus responsible for new cell infections would be difficult to distinguish within the total infectious viral load especially for rapidly growing viruses, such as CHIKV. Consequently, modeling occurrence of cell infections via total infectious viral load could result in misleading model parametrization. It is important to revise assumptions about viral infection dynamics as they could profoundly affect conclusions drawn from modeling *in vitro* virus dynamics under antiviral therapy, as the assays are often performed in adherent cell culture, as well as from modeling *in vivo* virus spread in organs and tissues.

In this study, we use viral dynamics modeling to numerically characterize the main determinants of ZIKV and CHIKV *in vitro* kinetics and to tease apart the effects of each determinant on the viral load. To inform the mathematical model, we measured temporal changes in the infectious viral titres and encapsulated genome abundances in a series of experiments reflective of different aspect of viral replication cycle in the extracellular milieu. To minimize the influence of immune responses on the CHIKV and ZIKV infection, we used a mammalian cell line (Vero) which is incapable of producing type I interferon in response to viral infections [46, 47]. Infection of Vero cells was carried out using two distinct amounts of input virus at multiplicity of infection (MOI, defined as the number of viral genomes that enter and effectively replicate in a cell) of 0.01 (hereafter referred to as low MOI) and 1 (hereafter referred to as high MOI) of infectious virus per cell. Using mathematical modeling, we compared CHIKV and ZIKV infection kinetics by allowing new infections to be facilitated via free extracellular infectious virus (hereafter referred to as ‘standard’ model) or via virus-producing cells (hereafter referred to as cell-to-cell transmission model). We show that because CHIKV-infected cells exhibit a much shorter eclipse phase and rapid accumulation of virus during the initial growth phase compared to rather long eclipse phase of ZIKV-infected cells and slow accumulation of virus over the infection course, the standard model fails to describe temporal CHIKV viral load data. The cell-to-cell transmission model, in which virus spread occurs via virus-producing cells, transpired to be significantly more descriptive of both CHIKV and ZIKV viral load time course data. Overall, we deliver the first comprehensive numerical characterization of *in vitro* CHIKV and ZIKV infections.

## Results

### Quantification of chikungunya and Zika loss of infectivity *c* and RNA genome stability *c*_rna_

To precisely calculate the degradation rates of infectious virus *c* and RNA genomes *c*_rna_, we experimentally measured stability of RNA genomes subjected to the physical conditions of the *in vitro* experiments. ZIKV and CHIKV stocks were incubated at 37°C for up to 72h in cell culture media and at time points 8h, 48h and 72h, RNA was extracted and quantified by qRT-PCR. By fitting equation (1) to RNA genome abundances (see Material and Methods for description of the fitting procedure), we determined that CHIKV RNA genome degradation over the course of 72h was negligible (Figure 1a) and for practical reasons was in the model (4) set to zero. ZIKV RNA genome degradation over the course of 72h was *c*_rna_ = 0.1h^−1^ (a half-life of 69.3h) (Figure 1b). Infectivity of both ZIKV and CHIKV were significantly reduced over time as determined by titration by plaque assay of infectious virus remaining in the solution (Figure 1a, 1b). By fitting equation (2) to viral titres (see Material and Methods for description of the fitting procedure), we determined the mean infectivity loss rate of CHIKV and ZIKV over the infection course to be *c* = 0.48h^−1^ and *c* = 0.72h^−1^, respectively, (a half-life of 14.4h and 9.6h, respectively). In conclusion, ZIKV loses infectivity more rapidly than CHIKV. Estimated viral decay kinetic parameters for both ZIKV and CHIKV with their 95% confidence intervals are summarized in Table 1 and Figure 1.

**Table 1:** Best-fit parameter values and 95% confidence intervals obtained from fitting equations (1) and (2) to total RNA genome abundances and viral titres, respectively, from the RNA genome degradation assays with the corresponding titre quantifications to asses infectivity of CHIKV (Figure 1a) and ZIKV (Figure 1b) over time.

| Description | Parameter | Value, 95% CI ($\text{h}^{-1}$) | |
| --- | --- | --- | --- |
|  |  | CHIKV | ZIKV |
| Infectious virus decay rate | $c$ | 0.05, [0.03, 0.06] | 0.072, [0.069, 0.076] |
| RNA genome decay rate | $c_{\text{rna}}$ | $2.2 \times 10^{-10}$, [$10^{-10}$, $3.7 \times 10^{-3}$] | 0.01, [0.0082, 0.011] |

**Figure 1:**
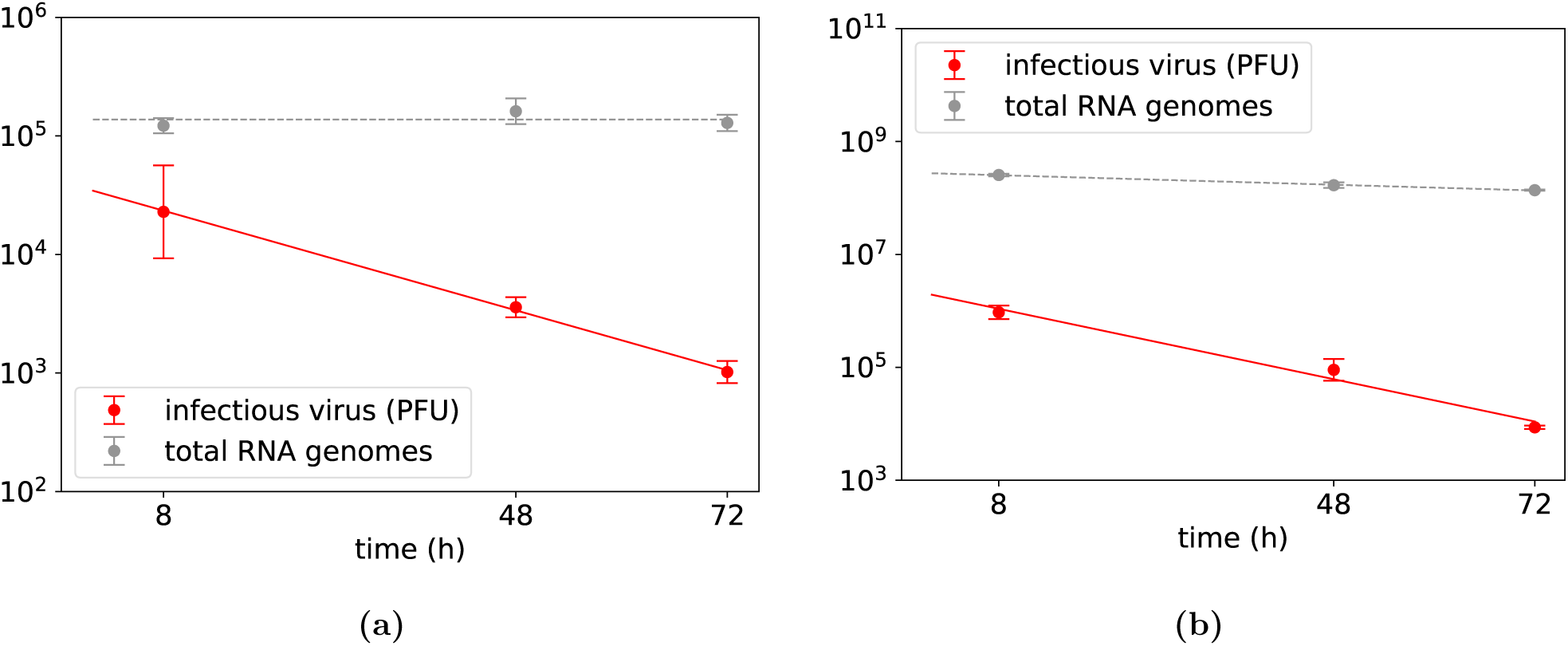
Cell-free experiment to determine stability of RNA genomes (grey dots) and loss of infectivity (red dots) of **(a)** CHIKV and **(b)** ZIKV subjected to the physical conditions of the *in vitro* experiments over 72h. The best fits of equations (1) and (2) describing the decay of RNA genomes and viral infectivity, respectively, are displayed as dashed and solid lines, respectively. Data are shown as the mean ± standard deviation. The best-fit parameter values and 95% confidence intervals are in Table 1.

### Model selection

For each virus, we used Approximate Bayesian Computation (ABC, Materials and Methods) to fit equations (4) (see also Figure 2 for biological description of the equations) to low and high MOI experimental datasets separately and simultaneously. For each virus and each MOI dataset, we quantified viral parameters within the model (4) for both viral transmission modes. To determine which of the two transmission models provides better description of the data, we performed model selection based on the calculation of posterior odds ratio (Materials and Methods). We found strong evidence for the cell-to-cell viral transmission model to describe CHIKV infection dynamics as the posterior odds ratio was equal to one in favour of cell-to-cell viral transmission model. For ZIKV, we also found evidence for the cell-to-cell viral transmission model to describe infection dynamics with posterior odds ratio equal to 0.74 for the cell-to-cell viral transmission model compared to 0.26 for the standard transmission model. Solutions of the cell-to-cell transmission model associated with parameter sets inferred from ABC provided good fits to both CHIKV and ZIKV time course datasets (Figures 3a and 4a, respectively). In contrast, solutions of the standard model associated with parameter sets inferred from ABC fit well ZIKV time course datasets (Figure 4b) and CHIKV time course datasets only when low and high MOI datasets were fit separately (results not shown) but did not fit well CHIKV time course datasets when low and high MOI datasets were fit simultaneously (Figure 3b).

**Figure 2:**
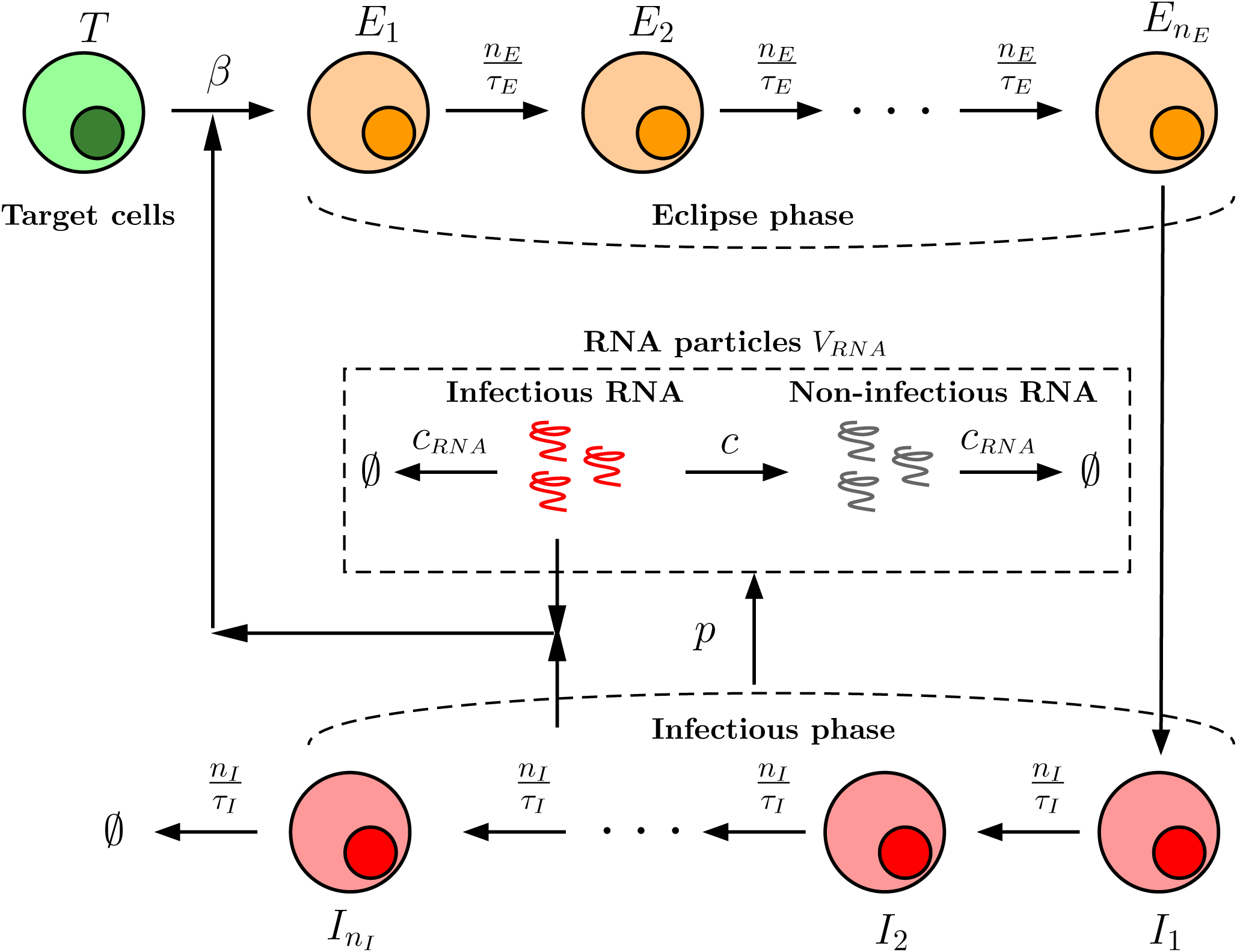
Graphical representation of the mathematical model (4) describing the *in vitro* viral kinetics. Susceptible cells (*T*) may get infected either by extracellular free virus entering susceptible cells at the rate (*β*_*V*_ *V*_pfu_) or when virus invades susceptible cells from virus-producing cells via cell-to-cell transmission at the rate (*β*_*C*_ *V*_pfu_). Upon successful virus infection, susceptible cells enter an eclipse phase which is divided into *n*_*E*_ sub-phases each of which last (*n*_*E*_*/τ*_*E*_) time units. Thus, the duration of eclipse phase is *τ*_*E*_ time units and 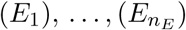 are cells in eclipse sub-phases. Only cells in the last sub-phase of eclipse phase 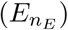 enter infectious phase in which they become virus-producing. Infectious phase is divided into *n*_*I*_ sub-phases each of which last (*n*_*I*_ */τ*_*I*_) time units. Thus, the duration of infectious phase is *τ*_*I*_ time units and 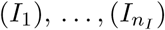 are cells in infectious sub-phases. Cells in any infectious sub-phase produce virus at the rate (*p*). Only cells in the last sub-phase of infectious phase 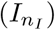 exit the system at the rate (*n*_*I*_ */τ*_*I*_). Infectious virus loses infectivity at the rate (*c*) and viral genomes lose stability at the rate (*c*_rna_).

**Figure 3:**
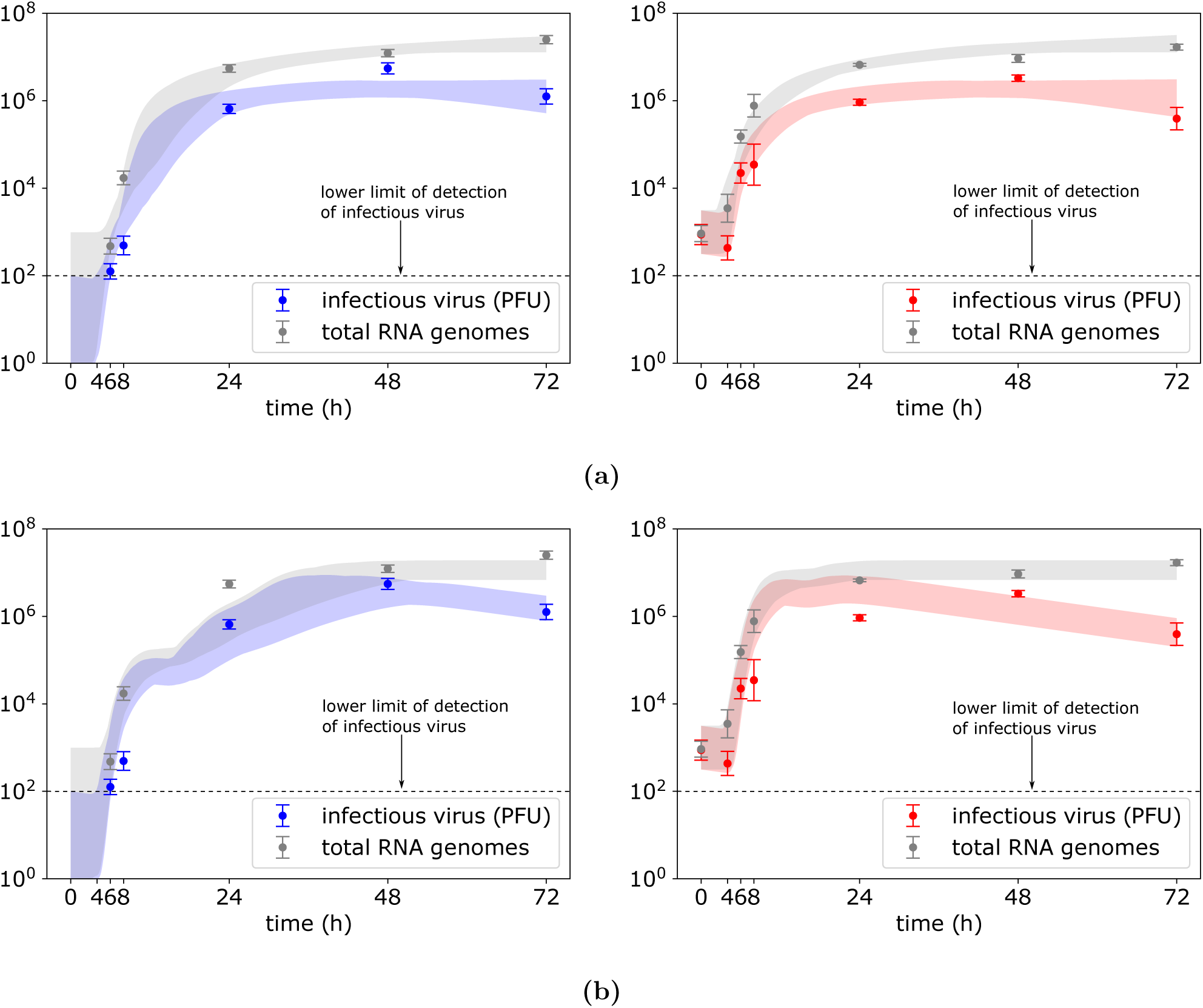
One thousand best ABC fits of the **(a)** standard **(b)** cell-to-cell transmission model to low (left panel) and high (right panel) MOI datasets depicted as filled areas around the time course CHIKV titres and total RNA genome abundances. Data are shown as the mean ± standard deviation and model was fit to low and high MOI datasets simultaneously.

**Figure 4:**
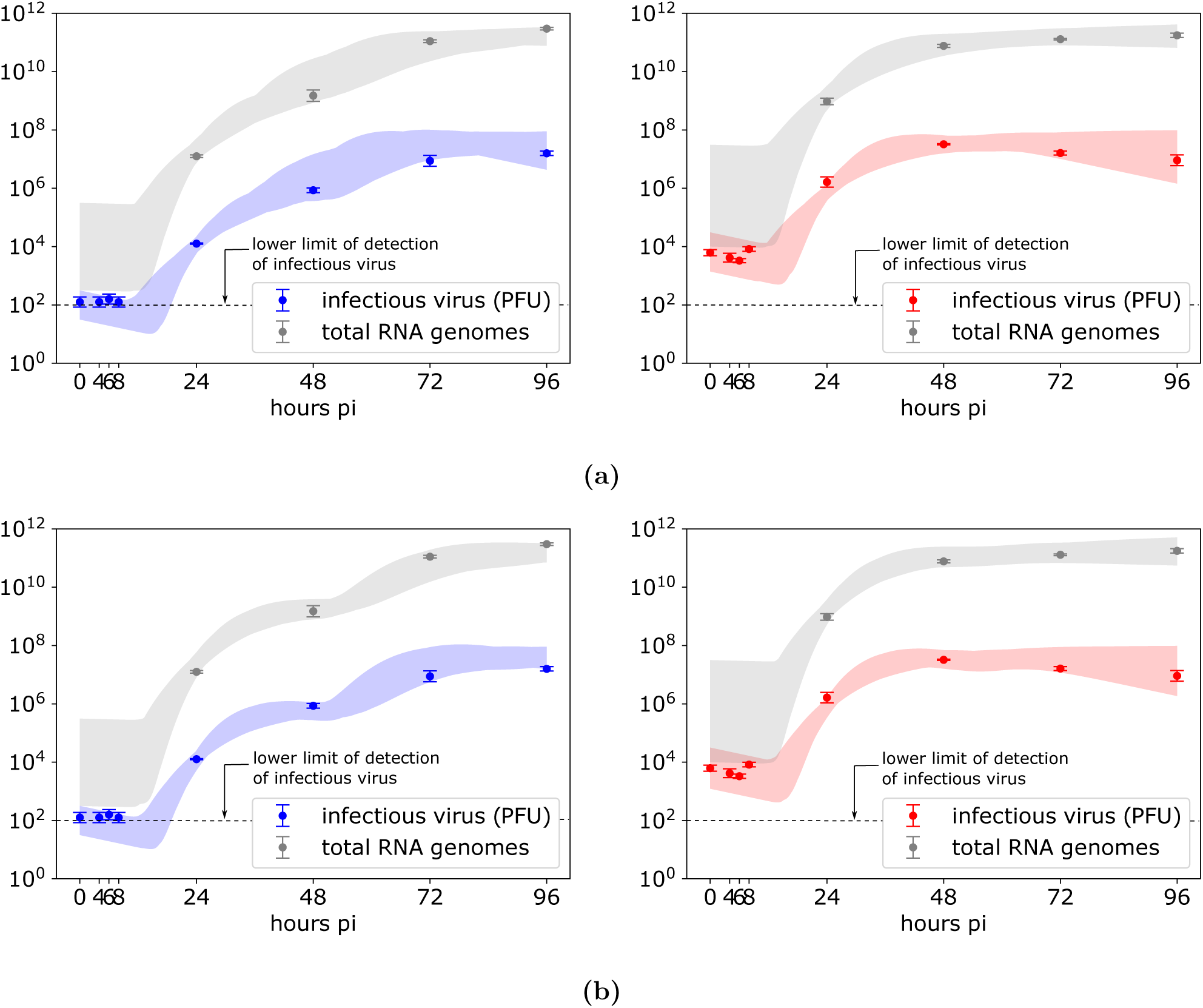
One thousand best ABC fits of the **(a)** standard **(b)** cell-to-cell transmission model to low (left panel) and high (right panel) MOI datasets depicted as filled areas around the time course ZIKV titres and total RNA genome abundances. Data are shown as the mean ± standard deviation and model was fit to low and high MOI datasets simultaneously.

Herein, we focus on the select model parameters, that is, eclipse phase *τ*_*E*_, viral genome production rate *p* and infectious virus to total RNA genomes ratio *α*. For the remaining model parameters, namely the number of compartments of eclipse and infectious phases *n*_*E*_ and *n*_*I*_, respectively, the mean lifespan of infected cells *τ*_*I*_ and the infection rate *β* the ABC converged on posterior distributions that were not significantly different from their uniform priors (results not shown). The mean, median and 95% credible intervals for all viral parameters for both models, both viruses and input MOI are listed in Tables 2, 3, 4 and 5.

**Table 2:**
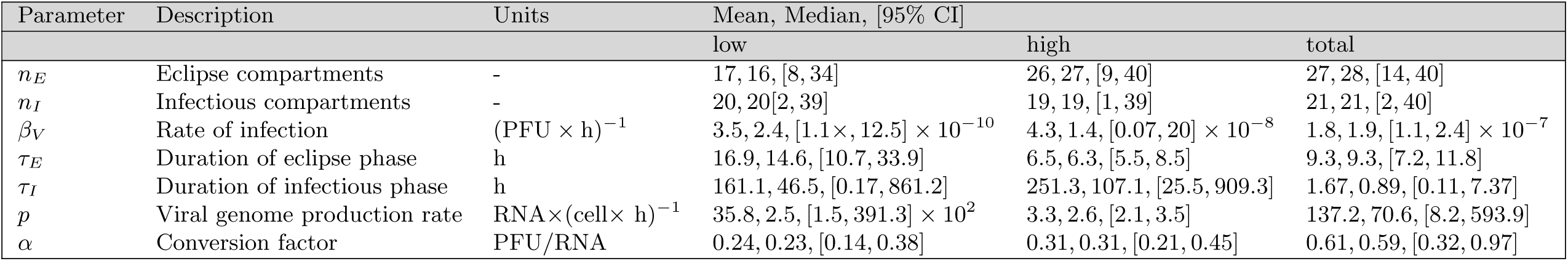
Mean, median and 95% credible intervals for viral parameters associated with ABC fits of the standard viral transmission model (equations (4)) to viral titres and RNA genome abundances obtained from low and high MOI yield assays of CHIKV infection.

**Table 3:**
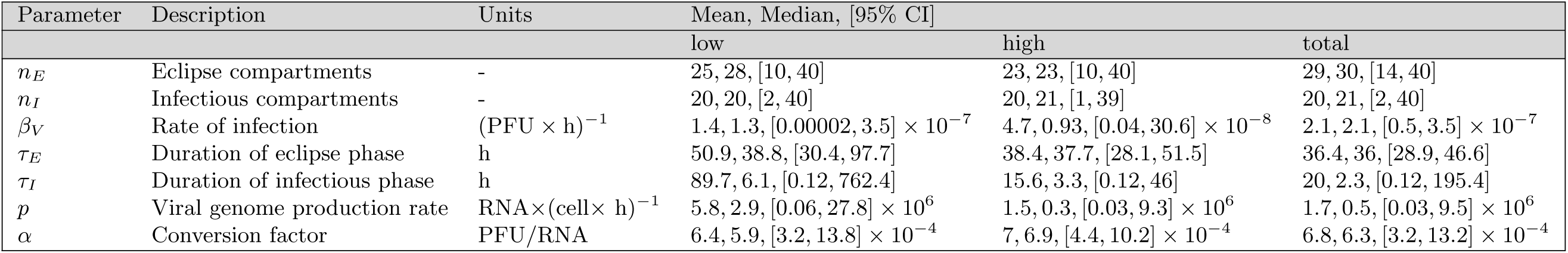
Mean, median and 95% credible intervals for viral parameters associated with ABC fits of the standard viral transmission model (equations (4)) to viral titres and RNA genome abundances obtained from low and high MOI yield assays of ZIKV infection.

**Table 4:** Mean, median and 95% credible intervals for viral parameters associated with ABC fits of the cell-to-cell viral transmission model (equations (4)) to viral titres and RNA genome abundances obtained from low and high MOI yield assays of CHIKV infection.

| Parameter | Description | Units | Mean, Median, [95% CI] |  |  |
| --- | --- | --- | --- | --- | --- |
|  |  |  | low | high | total |
| $n_E$ | Eclipse compartments | - | 21, 21, [5, 40] | 28, 29, [12, 40] | 25, 27, [8, 40] |
| $n_I$ | Infectious compartments | - | 20, 21, [2, 40] | 21, 22, [2, 40] | 21, 20, [2, 40] |
| $\beta_C$ | Rate of infection | $(\text{PFU} \times \text{h})^{-1}$ | 0.95, 0.04, $[4.8 \times 10^{-6}, 7.36]$ | 0.67, 0.005, $[1.54 \times 10^{-6}, 7.1]$ | 0.35, 0.007, $[7.59 \times 10^{-5}, 4.42]$ |
| $\tau_E$ | Duration of eclipse phase | h | 10.5, 7.5, [6.6, 34.3] | 6.1, 6.0, [5.3, 7.4] | 7.0, 6.7, [5.8, 9.1] |
| $\tau_I$ | Duration of infectious phase | h | 218.5, 98.6, [34.5, 876] | 244.7, 132.8, [0.3, 903.4] | 247.8, 134.5, [34.2, 875.2] |
| $p$ | Viral genome production rate | $\text{RNA} \times (\text{cell} \times \text{h})^{-1}$ | 1.89, 1.86, [1.5, 2.5] | 998.4, 2.4, [1.8, 8464.5] | 2.7, 2.0, [1.6, 2.7] |
| $\alpha$ | Conversion factor | $\text{PFU}/\text{RNA}$ | 0.24, 0.24, [0.16, 0.35] | 0.3, 0.3, [0.2, 0.4] | 0.3, 0.3, [0.2, 0.4] |

**Table 5:** Mean, median and 95% credible intervals for viral parameters associated with ABC fits of the cell-to-cell viral transmission model (equations (4)) to viral titres and RNA genome abundances obtained from low and high MOI yield assays of ZIKV infection.

| Parameter | Description | Units | Mean, Median, [95% CI] |  |  |
| --- | --- | --- | --- | --- | --- |
|  |  |  | low | high | total |
| $n_E$ | Eclipse compartments | - | 23, 23, [6, 39] | 22, 21, [7, 40] | 24, 24, [10, 40] |
| $n_I$ | Infectious compartments | - | 20, 20, [1, 39] | 20, 20, [2, 39] | 20, 20, [1, 39] |
| $\beta_C$ | Rate of infection | $(\text{PFU} \times \text{h})^{-1}$ | 0.035, 0.0015, $[6.3 \times 10^{-5}, 0.24]$ | 0.73, 0.008, $[1.63 \times 10^{-5}, 7.87]$ | 0.1, 0.003, $[6.56 \times 10^{-5}, 0.9]$ |
| $\tau_E$ | Duration of eclipse phase | h | 39.8, 37.3, [27.8, 74.4] | 37.9, 37.6, [25.3, 53.3] | 37.5, 36.4, [27.7, 55.4] |
| $\tau_I$ | Duration of infectious phase | h | 104.9, 10.3, [0.13, 845.7] | 30.7, 3.5, [0.1, 362.4] | 85.5, 7.0, [0.13, 790.5] |
| $p$ | Viral genome production rate | $\text{RNA} \times (\text{cell} \times \text{h})^{-1}$ | 1.6, 0.13, $[0.023, 10.8] \times 10^6$ | 1.4, 0.26, $[0.018, 8.2] \times 10^6$ | 1.2, 1.6, $[0.019, 8.2] \times 10^6$ |
| $\alpha$ | Conversion factor | $\text{PFU}/\text{RNA}$ | 7.1, 6.5, $[3.3, 14.1] \times 10^{-4}$ | 7.5, 7.4, $[4.4, 11.3] \times 10^{-4}$ | 7.4, 7.0, $[3.7, 12.7] \times 10^{-4}$ |

### Duration of eclipse phase *τ*_*E*_ of chikungunya- and Zika-infected cells is not exponential

Inference process under standard model yielded posterior distributions of *τ*_*E*_ with substantially different peaks and shapes for different initial experimental conditions (MOI) in the case of CHIKV infection (Figure 5a, left column). We estimated the median to be 14.6h for low MOI CHIKV infection, 6.3h for high MOI CHIKV infection and 9.3h if we fit the standard model to low and high MOI CHIKV datasets simultaneously. It is unlikely that differences in multiplicity of infection would promote such differences in posterior distributions of *τ*_*E*_ as the time for a virion to complete its replication cycle is biologically rather predetermined. Inference process under cell-to-cell transmission model converged to posterior distributions with consistent shapes and peaks for different initial experimental conditions (MOI) for both CHIKV and ZIKV infections (Figures 5b, 5d, left column) yielding medians between 6-7.5h and 36.4-39.8h, respectively. Interestingly, for ZIKV infection time course datasets, inference process under standard model yielded posterior distributions comparable to those under cell-to-cell transmission model (Figure 5c, left column) with the median between 36-38.8h across different initial viral input MOI. The mean, median and 95% credible intervals of the posterior distributions for the eclipse phase duration *τ*_*E*_ for each virus, each transmission model and each initial MOI are listed in Tables 2, 3, 4 and 5.

**Figure 5:**
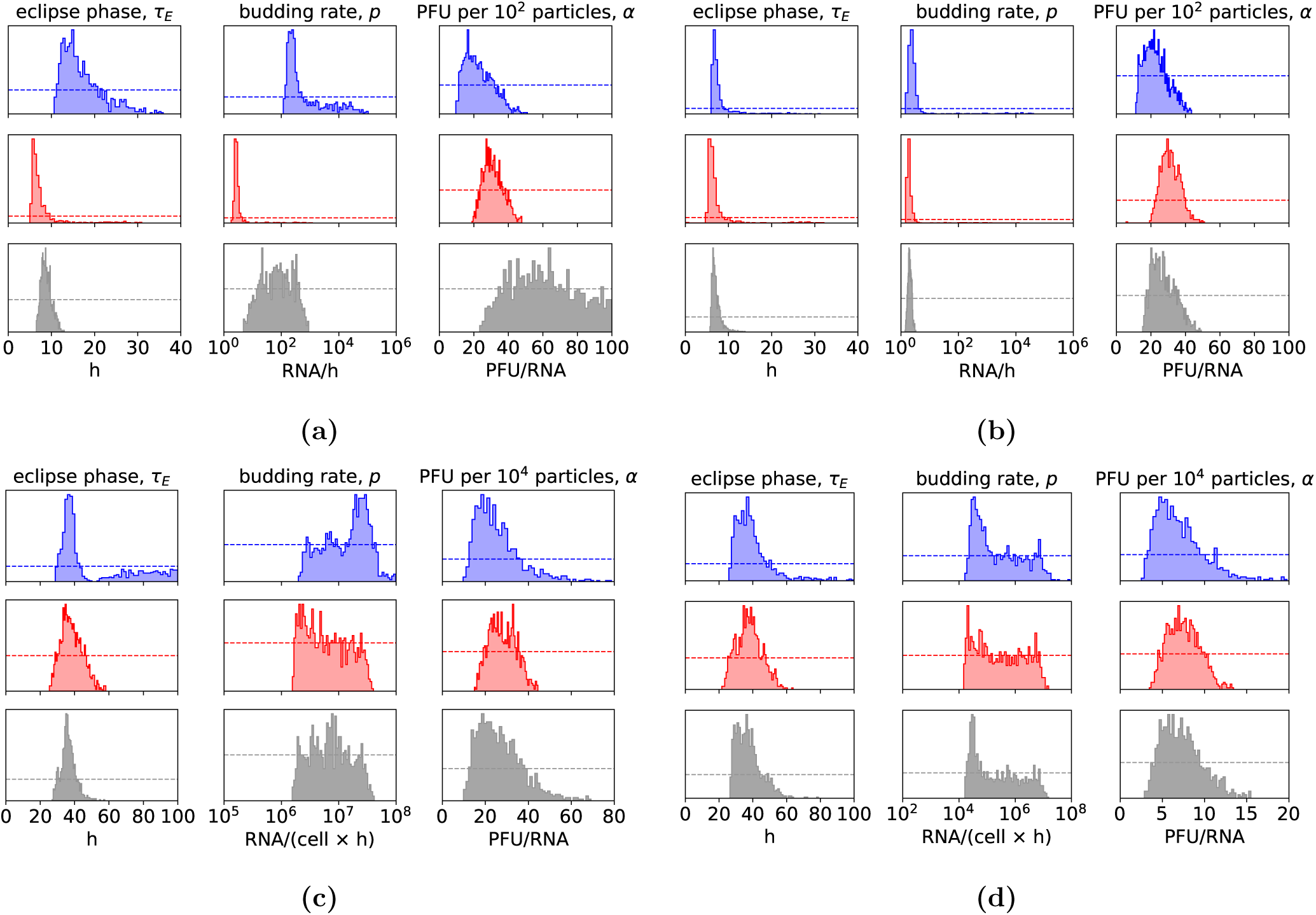
The posterior distributions of the select viral parameters obtained from ABC fits of the standard transmission model to **(a)** CHIKV and **(c)** ZIKV low (blue), high (red) and both low and high (grey) MOI kinetic data. The posterior distributions of the select viral parameters obtained from ABC fits of the cell-to-cell transmission model to **(b)** CHIKV and **(d)** ZIKV low (blue), high (red) and both low and high (grey) MOI kinetic data. The horizontal dashed lines indicate the initial prior (uniform) distribution from which the viral parameter values were sampled. The bounds imposed on viral parameters are in Material and Methods.

### Viral genome production rate *p* and infectious virus to total RNA genomes ratio *α* are substantially different for chikungunya and Zika viruses

Posterior distributions of the viral genome production rate *p* exhibited substantial differences in the peaks and shapes when the standard model was fit to low and high MOI CHIKV experimental datasets (Figures 5a, middle column). We estimated the median to be 250, 2.6, and 70.6, viral genomes released out of a cell per hour. These discrepancies disappeared when CHIKV dynamics was described by the cell-to-cell viral transmission model (Figure 5b) yielding median between 1.9-2.4 viral genomes per hour across different initial viral input MOI. Inference of ZIKV genome production rate under both standard and cell-to-cell transmission models yielded a bounded posterior distribution of *p* of which median varied between 0.13-2.9×10^6^ genomes released out of a cell per hour across different initial viral input MOI (Figures 5c and 5d, middle column). ZIKV-infected cells appeared to produce considerably more viral genomes but also significantly less infectious virus to total RNA genomes produced compared to CHIKV. For CHIKV, median of posterior distribution of *α* varied between 3 to 5 infectious viruses per ten RNA genomes produced whereas for ZIKV we obtained 5.9-7.4 infectious viruses per ten thousand RNA genomes produced (Figure 5, right column in each panel). The mean, median and 95% credible intervals of the posterior distributions for the viral genome production rate *p* and infectious virus to total RNA genomes ratio *α* for each virus, each transmission model and each initial MOI are listed in Tables 2, 3, 4 and 5.

### Viral parameters within cell-to-cell transmission model

We used the least-square fitting procedure described in Materials and Methods (Extraction of virus decay parameters) to precisely quantify the viral parameters by fitting the cell-to-cell transmission model (4) to low and high MOI datasets simultaneously. Because the number of compartments of the eclipse and infectious phases, *n*_*E*_ and *n*_*I*_, respectively, could not be inferred, we set *n*_*E*_ = *n*_*I*_ = 40 (as e.g. used to estimate Influenza A *in vitro* kinetic parameters in [29]) and fit equations (4) to low and high MOI CHIKV and ZIKV datasets. The estimated best-fit parameter values and 95% bootstrap confidence intervals are listed in Table 6 and associated dynamics of CHIKV and ZIKV infections are depicted in Figure 7. The infection rate *β*_*C*_ was found to be *β*_*C*_ = 4.2 × 10^−3^ and × 10^−4^ (cells×h)^−1^ for CHIKV and ZIKV infections, respectively. The duration of eclipse phase of CHIKV- and ZIKV-infected cells were found to be 6.4h and 29.4h, respectively. The mean lifespan of CHIKV- and ZIKV-producing cells were found to be 44.8h and 31.4h, respectively. Although the lifespan of virus-producing cells seems to be overestimated, especially in the case of CHIKV as it is highly cytopathic and promotes rapid cell death, such high values may be the result of post-peak virus clearance not having been captured in the data (Figures 7). As virus-producing cells undergo infection-induced death, additional data points capturing viral decay would reflect the phase when viral production becomes slower than viral clearance and possibly improve estimation of lifespan of virus-producing cells. The viral genome production rate *p* and infectious virus to total RNA genomes produced by a cell *α* were found to be significantly different for both viruses. While production rate of CHIKV genomes was estimated to be 2.4 genomes per cell per hour with the proportion of 18 infectious viruses per one hundred genomes, ZIKV genomes were being produced at the rate 3.3 × 10^4^ genomes per cell per hour with the proportion of 6.3 infectious viruses per ten thousand genomes produced (Table 6).

**Table 6:** Parameter values and 95% confidence intervals obtained from least-square fitting of equations (4) to viral titres and total RNA genome abundances from the high and low MOI CHIKV (Figures 7a and 7b) and ZIKV (Figures 7c and 7d) yield assays.

| Description | Parameter | Units | Value, [95% CI] |  |
| --- | --- | --- | --- | --- |
|  |  |  | CHIKV | ZIKV |
| Rate of infection | $\beta_C$ | $(\text{cell} \times \text{h})^{-1}$ | 4.2, $[0.03, 1154.5] \times 10^{-3}$ | 3.5, $[0.57, 24.6] \times 10^{-4}$ |
| Duration of eclipse phase | $\tau_E$ | h | 6.4, [5.6, 7.3] | 29.4, [25.4, 31.3] |
| Duration of infectious phase | $\tau_I$ | h | 44.8, [24.9, 87.6] | 31.4, [15.9, 169.4] |
| Viral genome production rate | $p$ | $\text{RNA} \times \text{h}^{-1}$ | 2.4, [1.5, 6.8] | 3.3, $[1.5, 7.8] \times 10^4$ |
| Conversion factor | $\alpha$ | $\text{PFU} \times \text{RNA}^{-1}$ | 0.18, [0.1, 0.26] | 6.3, $[2.2, 9.2] \times 10^{-4}$ |

## Discussion

The present study investigated the *in vitro* dynamics of chikungunya (CHIKV) and Zika (ZIKV) viruses whose time course of viral load data showed significantly different replication cycle kinetics. In particular, a longer replication cycle of ZIKV compared to that of CHIKV gave rise to qualitatively distinct viral dynamics which we studied by mathematical modeling to tease apart and quantify individual drivers within each virus-cell interactions. The dynamics of extracellular free virus was found not to be descriptive of either CHIKV or ZIKV infection dynamics. Therefore, we hypothesized that the rate at which cells were infected was not proportional to the total extracellular infectious virus but rather the number of virus-producing cells.

In modeling the viral kinetics, we were able to evaluate which of the two transmission models, that is the standard model in which viral transmission is facilitated by extracellular free virus or cell-to-cell transmission model in which viral transmission is facilitated by virus-producing cells, can explain empirical observations. Although the dynamics of virus-producing cells transpired to be significantly more explanatory of viral kinetic data, we cannot establish the exact mechanisms responsible. The cell-to-cell transmission term (−*β*_*C*_ *T* Σ_*j*_ *I*_*j*_) in the mathematical model (4) represents two physical and generally distinct biological processes; first, utilization of existing cell-to-cell contacts by the virus and second, exploitation of cell adhesion biology to deliberately establish contact between infected and uninfected target cells. Biologically, much remains unknown about the possibility of ZIKV and CHIKV spread via direct cell-to-cell interactions. To date, evidence for such spread for ZIKV does not exist. On the other hand, cell-to-cell-transmission of CHIKV has been previously suggested to describe the resistance of CHIKV mutants to antibody-dependent neutralization [36]. The authors suggested that presumably cell-to-cell transmission occurs when virus budding occurs near a cell junction and when the virus can recognize the viral receptors on the neighboring cell. It is important to note that other ways of transmission, which may resemble cell-to-cell transmission in ‘protecting’ the virus from the extracellular space exist, and have not been taken into account in the latter study. For example, it is becoming increasingly evident that viruses hijack cellular machinery to be transmitted through extracellular vesicles (such as exosomes) in order to escape antibody and immune responses and mediating further infection [48–51]. Indeed, ZIKV transmission has been shown to be mediated by exosomes in cortical neurons [52]. In a similar manner, CHIKV was shown to trigger apoptosis and ‘hide’ in apoptotic blebs, which were then able to infect cells otherwise refractory to CHIKV infection [49]. Although direct cell-to-cell viral transmission remains to be experimentally explored and demonstrated for CHIKV and ZIKV, we showed that mathematical model in which virus spread is proportional to virus-producing cells is able to explain experimental data more accurately. Despite that the spatial component of virus infection dynamics is not taken into account by either of the two models, a virus is most likely to infect neighboring cells following budding, especially in more static environments such as cell culture without mixing, or possibly in tissue compartments *in vivo*. This may explain why the cell-to-cell transmission model is favored in this study, as the extracellular free virus model, which assumes that progeny virus is likely to infect any cell, irrespective of the distance from the infected cell.

It is important to note that most empirical data used for modeling tend to be generated under a single growth condition. We conscientiously chose to perform and analyze both low and high MOI growth, individually and combined, to further test the validity of each model. Had only one growth condition been selected, we would not have identified that the standard model failed. These results argue for the inclusion of more diverse experimental sets in model selection and development. We could argue that the extreme differences between the inferred posterior distributions of CHIKV viral parameter values under free-virus transmission may have been the result of MOI-dependent cellular response to the presence of the virus throughout the infection course. Another possible explanation for the reported discrepancies is superinfection. However, CHIKV superinfection is not well supported since it has recently been shown that prior CHIKV infection of BHK cells (which are also interferon-incompetent) inhibits re-infection of already infected cells by a challenge CHIKV [53]. Modeling CHIKV *in vitro* dynamics thus presents a challenge and requires further investigation.

Statistical model comparison provided more support for cell-to-cell over the standard viral transmission model. Nevertheless, this does not imply that cell-to-cell transmission model is the correct model to be used to model CHIKV *in vitro* dynamics. To possibly test the hypothesis that standard transmission model is indeed descriptive of CHIKV kinetics, infectious and total RNA genomes would have to be measured in a timely manner in-between time points 8h and 24h to capture two-phase increase of virus, particularly at low MOI. Insufficient data may have not provided enough information about the viral dynamics to the mathematical model. Genome quantification of intracellular virus would provide evidence for differential accumulation of virus within cells during low and high MOI infection. Another, although indirect, evidence to support the standard model would include timely measures of cell death as their accumulation would reflect removal of short-lived virus producing cells with a large viral burst size from the system. Interestingly, both standard and cell-to-cell transmission models were able to describe ZIKV in vitro kinetics. Although there was stronger statistical preference for the latter, inference process yielded comparable posterior distributions of viral parameter values for both standard and cell-to-cell transmission models. Thus, we conclude that CHIKV as a model virus with fast replication cycle may exhibit MOI-dependent phenotypes.

Overall, this study showed that the mathematical model in which the spread of an infection is described by cell-to-cell viral transmission can more accurately describe the *in vitro* dynamics of CHIKV and ZIKV infections than the standard model in which the spread of an infection is mediated via free extracellular virus. By modeling viral load datasets reflective of the virus kinetics at low and high MOI, we quantified the rates of different processes within the CHIKV- and ZIKV-cell interactions. Although we could not directly identify and quantify specific mechanisms, differences in the time scales of viral replication cycle may play an important role in identifying the model of better predictive power. This could have a significant impact on the development of models for viral control as the predictive ability of a chosen model to reflect and meaningfully interpret viral data under the influence of an external intervention, such as antiviral treatment, would be skewed. Identifying descriptive models and confronting them with diverse experimental datasets is essential to the development of therapies that prevent or treat CHIKV, ZIKV, and other infections.

## Materials and Methods

### Cells

Vero, HEK-293T and BHK cells were maintained in Dulbecco’s modified Eagles medium (DMEM), supplemented with 10% fetal calf serum (FCS) and 1% penicillin/streptomycin (P/S; Thermo Fisher) in a humidified atmosphere at 37°C with 5% CO2. U4.4 cells (derived from Aedes albopictus) were grown in Leibovitz’s L-15 medium with 10% FCS, supplemented with 1% P/S, 1% non-essential amino acids (Sigma) and 1% tryptose phosphate (Sigma) at 28°C.

### Viruses

The chikungunya virus (CHIKV) stock was generated from a Caribbean infectious clone described elsewhere [54]. After linearization with *NotI* restriction enzyme (Thermo Fisher), RNA was generated by *in vitro* transcription (IVT) with SP6 mMESSAGE mMACHINE kit (Invitrogen) and transfected into BHK with lipofectamin 2000 (invitrogen). The Zika virus (ZIKV) used for this study is the prototype african MR-766 strain derived from an infectious clone described elsewhere [55]. ZIKV was rescued by transfection of the infectious clone in semi-confluent HEK-293T cells using TransIT-LT1 transfection reagent (Mirus Bio). For both viruses, virus stocks used in this study were generated by infection of Vero cells for amplification, titred by plaque assay and frozen at −80°C prior to use.

### Plaque assay

Viral titration was performed on Vero cells plated 1 day prior to infection on 24 well plates. Ten fold dilutions were performed in DMEM alone and transferred onto Vero cells for 1 hour to allow infection before adding DMEM with 2% FCS, 1% P/S and 0.8% agarose. Plaque assays were fixed with 4% formalin (Sigma) 3 days post infection (p.i.) (CHIKV) or 4 days p.i. (ZIKV) and plaques were manually counted.

### Growth curves

Cells were plated in 12 well plates at 80-90% confluence one day before infection. At day 0, virus was diluted in 300 or 200 *µ*l PBS to obtain a multiplicity of infection (MOI) of 1 PFU per cell (high MOI) or 0.01 PFU per cell (low MOI). After 1 hour, the viral solution was removed, cells were washed three times with PBS and new media supplemented with 2% FCS was added. At each time point 0h, 4h, 8h, 24h, 48h and 72h for CHIKV and 0h, 4h, 6h, 8h, 24h, 48h, 72h, and 96h for ZIKV infection 60 *µ*l and 5 *µ*l were separately aliquoted and frozen for further titration (as described above) and RT qPCR. 65 *µ*l of fresh media was added on top of cells to replace the taken volume. Each growth curve was done in triplicates.

### RT qPCR

As described in [56], cell supernatants were heated 5 minutes at 60°C for viral inactivation. Quantitative RT-PCR was then performed with TaqMan RNA-to-Ct One-step RT-PCR kit (Applied Biosystems) using the following cycling conditions: 20 minutes at 50 C, 10 minutes at 95 C, 40 cycles of 95 C for 15 seconds, followed by 60 C for 1 minute). The primer and probe sets used for each virus are shown in Table 7. RNA copy number was derived from a standard curve generated using reactions containing 10-fold dilutions of known amounts of in vitro generated RNA transcripts Each reaction contained a scale of diluted IVT to calculate RNA copy number. The CHIKV RT-PCR amplifies a 152 nucleotide-region spanning the 5’ UTR and NSP1. The ZIKV primers bind to and amplify a 77 nucleotide region in the 5’ end of the ZIKV genome (position 1192-1268).

**Table 7:**
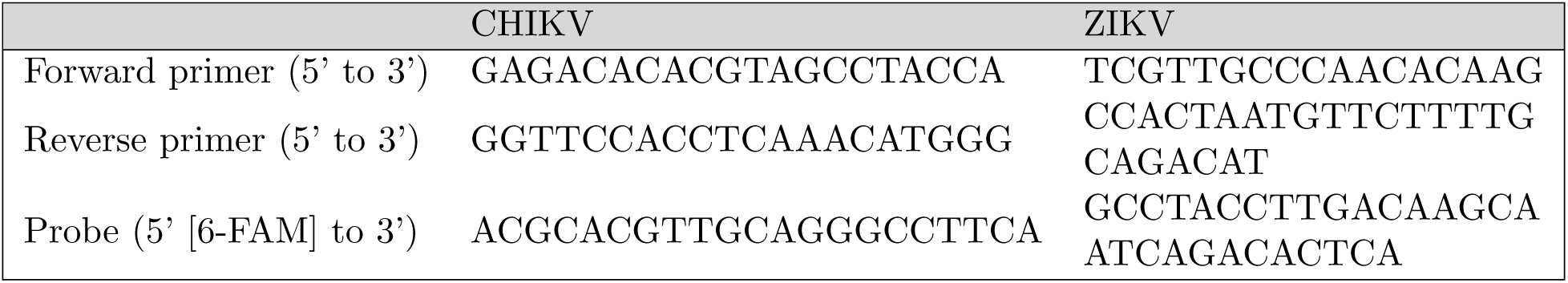
The primer and probe sets used for CHIKV and ZIKV.

### Extraction of virus decay parameters

To identify viral decay parameters, *c*_rna_ and *c*, we assumed that the loss of RNA genomes and infectious virus proceeds in an exponential (or log-linear) manner over time. Therefore, the loss of RNA genomes *V*_rna_(*t*) and infectious virus *V*_pfu_(*t*) can be expressed as

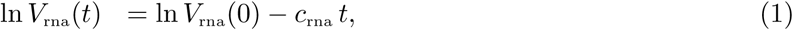

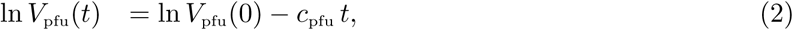

where *c*_rna_ and *c*_pfu_ (measured in h^−1^) are the decay rates of RNA genomes and virus infectivity, respectively, as in equations (4), and ln *V*_rna_(0) and ln *V*_pfu_(0) are natural logarithms of the initial states of RNA genomes and infectious virus at *t* = 0h.

To obtain estimates of viral decay parameters, we minimized the sum of squared errors (SSE) between the measured data *D*(*t*_*i*_) and the model solution *V* (*t*_*i*_) at each measured time point *t*_*i*_ and for each measured replicate *j* given as

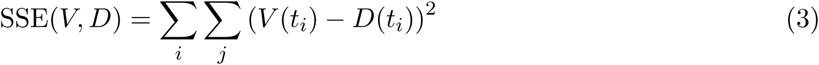

across parameter ranges using the Python function `scipy.optimize.least_squares` for performing bound-constrained optimization on variables. To provide 95% confidence intervals for each estimated best-fit parameter, we fit equations (1) and (2) to 3000 bootstrap replicates of each data set, the detailed description of which can be found in [59].

### Mathematical model

The cell-free, low and high MOI time course datasets were numerically simulated using a collection of ordinary differential equations, in which susceptible target cells (*T*) become infected by infectious virus ((*V*_pfu_), measured in plague forming units (PFU)) or virus-producing cells 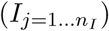 at the infection rate (*β*_*V*_), (measured in (PFU × h)^−1^) or (*β*_*I*_), (measured in (cell × h)^−1^), respectively. The rate of cell infection by infectious virus depends on the concentration of free extra-cellular infectious virus (*V*_pfu_), but not released total virus ((*V*_rna_), measured as total RNA genomes (RNA)). Upon successful infection, target cells enter an eclipse phase (the time between virus entry into the cell to the beginning of viral release out of the cell), separated into (*n*_*E*_) stages. Eclipse cells 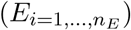 remain in each stage *i* = 1, …, *n*_*E*_ for an exponentially-distributed time of equal average length (*τ*_*E*_*/n*_*E*_). Only eclipse cells in the last compartment 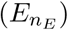 are allowed to transition into the infectious state and begin producing viral genomes. Infectious phase (the amount of time between the beginning of viral release out of a cell until the cell undergoes cell death or is removed from the state of being infectious by other mechanisms) is separated into (*n*_*I*_) stages, and infectious cells 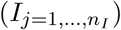 spend an exponentially-distributed time of equal average length (*τ*_*E*_*/n*_*E*_) in each stage before infectious cells in the last stage (*n*_*I*_) are removed from the system. It is unrealistic to impose the assumption on the cells to spend an exponentially distributed amount of time in the eclipse or infectious phases (here equivalent to *n*_*E*_ = *n*_*I*_ = 1) as it would allow cells to initiate viral production immediately upon infection, stop viral production immediately after it is initiated, and produce virus indefinitely [19, 18, 30, 31, 57]. Therefore, we subdivided eclipse and infectious phases into *n*_*E*_ and *n*_*I*_ compartments, such that these times are Erlang distributed with means *τ*_*E*_ and *τ*_*I*_, respectively. Erlang distribution is a special case of Gamma distribution of which the shape is dictated by *n*_*E*_ and *n*_*I*_ and vary from an exponential (= 1) to a normal-like (≫ 1) (Figure 6). Infectious cells in all stages can produce viral genomes at the rate ((*p*), measured in RNA×(cell × h)^−1^), the proportion of which ((*α*), measured in PFU/RNA) translates into infectious virus. Viral particles degrade at the rate ((*c*_rna_), measured in h^−1^) and infectious virus loses infectivity at the rate ((*c*), measured in h^−1^). The viral dynamics model is illustrated in Figure 2 and comprises of the following collection of ordinary differential equations:

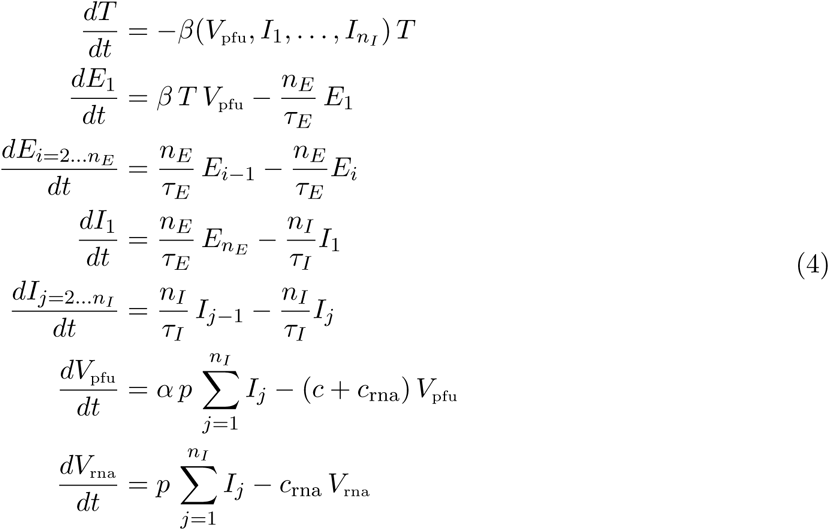

where

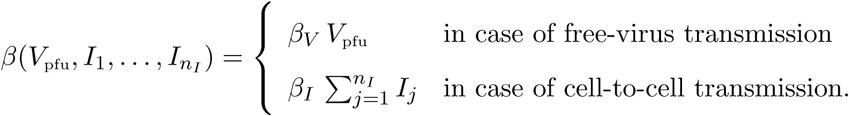

**Figure 6:**
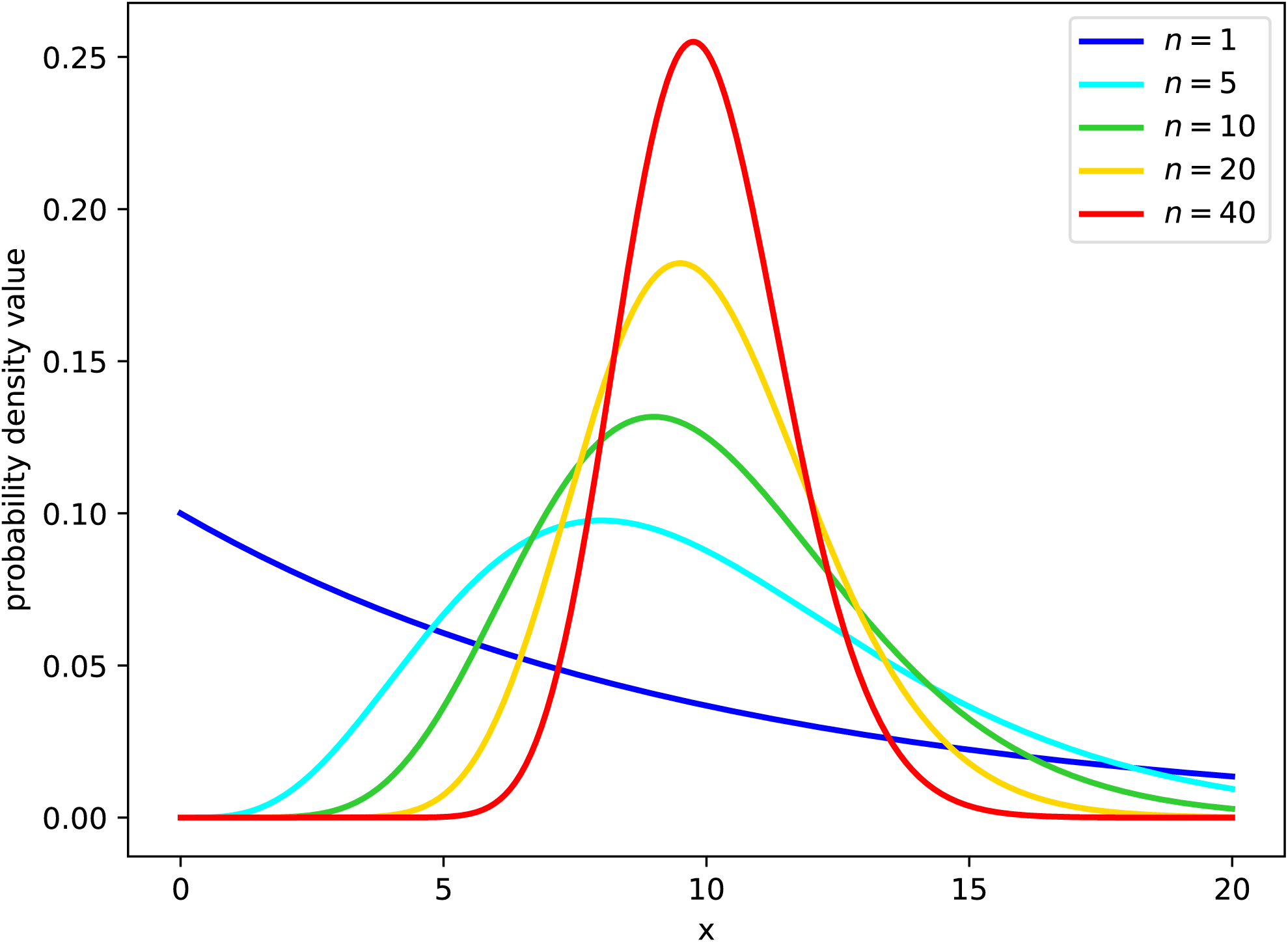
Probability density (y-axis) that a cell spends *x* hours in the (eclipse or infectious) phase. As the Erlang shape parameter (*n*_*E*_ or *n*_*I*_ in the model (4) for the eclipse and virus-producing phases, respectively) is increased, the distribution of the phase duration shifts from an exponential (*n* = 1), to a fat-tailed (1 < *n* < 10), to a normal-like (*n* ≫ 10) distribution. In these graphs, the mean time spent by cells in the phase (*τ*_*E*_ or *τ*_*I*_ in the model (4), respectively) is fixed (set to 10h, chosen arbitrarily) as the shape parameter (*n*_*E*_ or *n*_*I*_) is varied.

**Figure 7:**
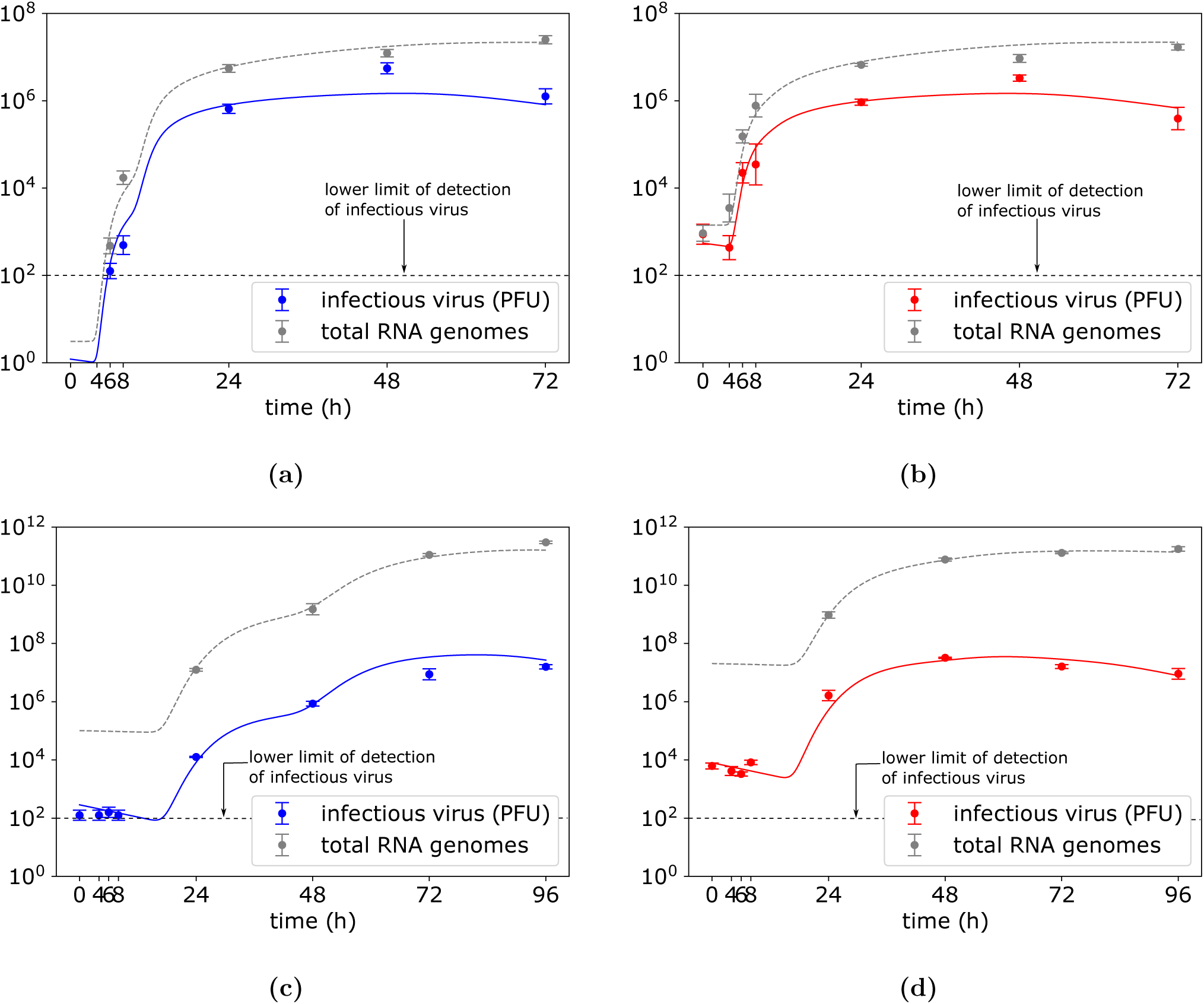
CHIKV and ZIKV kinetics that corresponds to the best-fit parameters obtained from the least-square fitting of the cell-to-cell viral transmission model (4) to **(a)** low and **(b)** high CHIKV MOI dataset, and **(c)** low and **(d)** high ZIKV MOI dataset. Data are shown as the mean ± standard deviation. The best-fit parameter values dictating CHIKV kinetics are in Table 6.

The experiments to obtain viral load data at different time points began with overlaying the virus supernatant on susceptible cells followed by a one hour cultivation to allow cell infection. The supernatant was then removed and cells were thoroughly washed off the remaining virus that did not enter the cells. The proportion of susceptible cells that became infected was governed by the multiplicity of infection (i.e., the ratio of infectious virus in the inoculum to the total number of susceptible cells) and was assumed to follow Poisson distribution as follows:

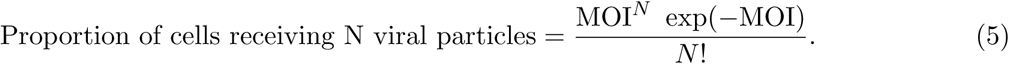

We further simplify the process by allowing only eclipse cells in their first stage of eclipse phase, *E*_1_, to have received the virus. The fraction of *E*_1_ cells which received one or more viruses is equivalent to the total proportion of cells excluding those which did not receive any virus

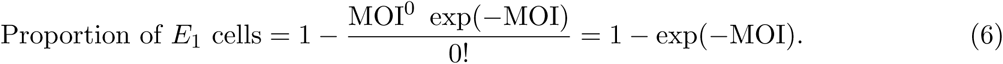

Thus, the initial conditions are *T* (*t* = 0) = *T*_0_ × exp(−MOI), *E*_1_(*t* = 0) = *T*_0_ × (1 − exp(−MOI)), 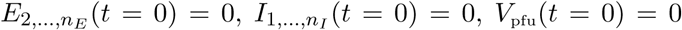, and *V*_rna_(*t* = 0) = 0, where *T*_0_ = 2 × 10^5^ susceptible cells seeded in each well.

### Parameter inference

We used Approximate Bayesian Computation (ABC) rejection approach to infer viral parameters and perform model selection. We simulated large numbers of datasets using parametrisations sampled from a log-uniform prior probability distribution for each model parameter. Specifically, the ranges over which we varied both CHIKV and ZIKV parameters were *n*_*E*_ ∼ *U* (1, 40), *n*_*I*_ ∼ *U* (1, 40), log_10_ *β*_*V*_ ∼ *U* (−10, *c/T*_0_), log_10_ *β*_*C*_ ∼ *U* (−6, −1), log_10_ *τ*_*E*_ ∼ *U* (−1, 2) and log_10_ *τ*_*I*_ ∼ *U* (−1, 3). CHIKV-specific parameter ranges were within log_10_ *p* ∼ *U* (0, 6), log_10_ *α* ∼ *U* (−2, 0) and ZIKV-specific parameter ranges were within log_10_ *p* ∼ *U* (2, 8), log_10_ *α* ∼ *U* (−5, 0). At the time point *t* = 0h, the extracellular virus was either undetectable or some residual virus was detected due to insufficient washing of cells. Thus, we keep the initial residual extracellular viral loads as free parameters and do not allow the residual virus to engage in the dynamics (residual infectious virus and RNA genomes are only allow to decay at the rates *c* and *c*_rna_, respectively). The initial CHIKV residual inputs were varied within log_10_ *V*_pfu_(0) ∼ *U* (0, 2), log_10_ *V*_rna_(0) ∼ *U* (0, 3) for low MOI infection and log_10_ *V*_pfu_(0) ∼ *U* (2.5, 3.5) and log_10_ *V*_rna_(0) ∼ *U* (2.5, 3.5) for high MOI infection. The initial ZIKV residual inputs were varied within log_10_ *V*_pfu_(0) ∼ *U* (1.5, 2.5), log_10_ *V*_rna_(0) ∼ *U* (2.5, 5.5) for low MOI infection and log_10_ *V*_pfu_(0) ∼ *U* (3, 4.5) and log_10_ *V*_rna_(0) ∼ *U* (4, 7.5) for high MOI infection. We numerically solved the system (4) using the Python function `scipy.integrate.odeint` and simulated data *V*_pfu_ and *V*_rna_ were then compared with the mean of measured data *D*_pfu_ and *D*_rna_ by calculating Euclidean distance

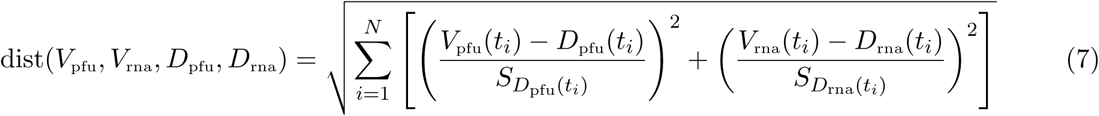

for measured times {*t_i_*|*i* = 1, …, *N*} where 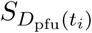 and 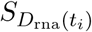 are the standard deviations of the measured viral titres and RNA genome abundances at the time point *t*_*i*_. The parametrizations of all simulated datasets were sorted with respect to the distance (7) in an ascending manner and the first one thousand parametrizations were accepted. The posterior density distributions were then constructed from the pool of accepted parametrisations.

### Model selection

We evaluate the evidence provided by the data in favour of cell-to-cell transmission model over the standard model by computing posterior odds as a summary of such evidence. Practically, for each of the models we find the largest distance (7) at which a parameter set was accepted. Using the smaller of these two distances, we can then determine for each model the number of parameter sets that would be accepted at this threshold. The posterior odds are then the fraction of all parameter sets accepted at this threshold contributed by each model.

### Biological constraints

We impose realistic biological constraints on the viral parameters whenever the spread of infection is modelled by free-virus transmission. Since infectious virus is cleared at a rate *c*, then its mean lifetime is 1*/c*. Therefore, the mean number of cells infectious virus infects during its lifetime is *β*_*V*_ *T*_0_*/c*. We require that inferred viral parameters satisfy *β*_*V*_ *T*_0_*/c ≤* 1 and thus, on average, infectious virus can infect at most one cell.

Since we initiate equations (4) at time 0h assuming a portion of cells already in the eclipse phase, any combination of viral parameters will result in viral growth. To ensure realistic parametrization of equations (4), we required the basic reproduction number *R*_0_, defined as the number of secondary infected cells that will be infected by a single infectious cell in a completely susceptible population is at least one, to satisfy *R*_0_ = *β*_*V*_ *T*_0_ *τ*_*I*_ *α p/c ≥* 1. *R*_0_ is here a product of the mean amount of infectious virus produced during the lifetime of an infected cell (*α p τ*_*I*_) and the mean number of cells infected per infectious virus *β*_*V*_ *T*_0_*/c* [58].

## Acknowledgements

We thank Carolina B. López and the members of Vignuzzi lab for discussions. This work was funded by the DARPA INTERCEPT program managed by Dr. Jim Gimlett and administered though DARPA Cooperative Agreement #HR0011-17-2-0023 (the content of the information does not necessarily reflect the position or the policy of the U.S. government, and no official endorsement should be inferred).

## References

[1] Robinson MC. An epidemic of virus disease in Southern Province, Tanganyika territory, in 19521953. 1955. Transactions of the Royal Society of Tropical Medicine and Hygiene 49(1):28– 32.

[2] Schuffenecker I, Iteman I, Michault A, Murri S, Frangeul L, Vaney MC, Lavenir R, Pardigon N, Reynes JM, Pettinelli F, Biscornet L. 2006. Genome microevolution of chikungunya viruses causing the Indian Ocean outbreak. PLOS Medicine 3(7):e263.

[3] Chen R, Puri V, Fedorova N, Lin D, Hari KL, Jain R, Rodas JD, Das SR, Shabman RS, Weaver SC.2016. Comprehensive genome scale phylogenetic study provides new insights on the global expansion of chikungunya virus. Journal of Virology 90(23):10600–11.

[4] Dick GW, Kitchen SF, Haddow AJ. Zikavirus (I). 1952. Isolations and serological specificity. Transactions of the Royal Society of Tropical Medicine and Hygiene 46(5):509–20.

[5] Lanciotti RS, Kosoy OL, Laven JJ, Velez JO, Lambert AJ, Johnson AJ, Stanfield SM, Duffy MR. 2008. Genetic and serologic properties of Zika virus associated with an epidemic, Yap State, Micronesia, 2007. Emerging Infectious Diseases 14(8):1232.

[6] Oehler E, Watrin L, Larre P, Leparc-Goffart I, Lastere S, Valour F, Baudouin L, Mallet HP, Musso D, Ghawche F. 2014. Zika virus infection complicated by Guillain-Barre syndromecase report, French Polynesia, December 2013. Eurosurveillance 19(9):20720.

[7] Dyer O. Zika virus spreads across Americas as concerns mount over birth defects. 2015. BMJ 351:h6983.

[8] Parra B, Lizarazo J, Jiménez-Arango JA, Zea-Vera AF, González-Manrique G, Vargas J, Angarita JA, Zuñiga G, Lopez-Gonzalez R, Beltran CL, Rizcala KH. 2016. GuillainBarré syndrome associated with Zika virus infection in Colombia. New England Journal of Medicine 375(16):1513– 23.

[9] Mlakar J, Korva M, Tul N, Popović M, Poljšak-Prijatelj M, Mraz J, Kolenc M, Resman Rus K, Vesnaver Vipotnik T, Fabjan Vodušek V, Vizjak A. 2016. Zika virus associated with microcephaly. New England Journal of Medicine 374(10):951–8.

[10] European Centre for Disease Prevention and Control. Rapid risk assessment: Zika virus epidemic in the Americas: potential association with microcephaly and Guillain-Barré syndrome. 2015.

[11] Ioos S, Mallet HP, Goffart IL, Gauthier V, Cardoso T, Herida M. Current Zika virus epidemiology and recent epidemics. 2014. Medecine et Maladies Infectieuses 44(7):302–7.

[12] Weger-Lucarelli J, Ruckert C, Chotiwan N, Nguyen C, Luna SM, Fauver JR, Foy BD, Perera R, Black WC, Kading RC, Ebel GD. 2016. Vector competence of American mosquitoes for three strains of Zika virus. PLOS Neglected Tropical Diseases 10(10):e0005101.

[13] Moser LA, Boylan BT, Moreira FR, Myers LJ, Svenson EL, Fedorova NB, Pickett BE, Bernard KA. Growth and adaptation of Zika virus in mammalian and mosquito cells. 2018. PLoS Neglected Tropical Diseases 12(11):e0006880.

[14] Perelson AS, Ribeiro RM. 2013. Modeling the within-host dynamics of HIV infection. BMC Biology 11(1):96.

[15] Wu H, Zhu H, Miao H, Perelson AS. Parameter identifiability and estimation of HIV/AIDS dynamic models. 2008. Bulletin of Mathematical Biology 70(3):785–99.

[16] Verotta D. 2005. Models and estimation methods for clinical HIV-1 data. Journal of Computational and Applied Mathematics 184(1):275–300.

[17] Miao H, Dykes C, Demeter LM, Cavenaugh J, Park SY, Perelson AS, Wu H. 2008. Modeling and estimation of kinetic parameters and replicative fitness of HIV-1 from flow-cytometry-based growth competition experiments. Bulletin of Mathematical Biology 70(6):1749–71.

[18] Kakizoe Y, Nakaoka S, Beauchemin CA, Morita S, Mori H, Igarashi T, Aihara K, Miura T, Iwami S. 2015. A method to determine the duration of the eclipse phase for in vitro infection with a highly pathogenic SHIV strain. Scientific Reports 5:10371.

[19] Beauchemin CA, Miura T, Iwami S. 2017. Duration of SHIV production by infected cells is not exponentially distributed: Implications for estimates of infection parameters and antiviral efficacy. Scientific Reports 7:42765.

[20] Chatterjee A, Guedj J, Perelson AS. 2012. Mathematical modeling of HCV infection: what can it teach us in the era of direct antiviral agents?. Antiviral Therapy 17(6 0 0):1171.

[21] Aston P. 2018. A new model for the dynamics of hepatitis C infection: Derivation, analysis and implications. Viruses 10(4):195.

[22] Rihan FA, Sheek-Hussein M, Tridane A, Yafia R. 2017. Dynamics of hepatitis C virus infection: mathematical modeling and parameter estimation. Mathematical Modeling of Natural Phenomena 12(5):33–47.

[23] Arthur JG, Tran HT, Aston P. Feasibility of parameter estimation in hepatitis C viral dynamics models. 2017. Journal of Inverse and Ill-Posed Problems 25(1):69–80.

[24] Perelson AS, Ribeiro RM. 2008. Estimating drug efficacy and viral dynamic parameters: HIV and HCV. Statistics in medicine 27(23):4647–57.

[25] Krakauer DC, Komarova NL. 2003. Levels of selection in positivestrand virus dynamics. Journal of evolutionary biology 16(1):64–73.

[26] Regoes RR, Crotty S, Antia R, Tanaka MM. 2005. Optimal replication of poliovirus within cells. The American Naturalist 165(3):364–73.

[27] Schulte MB, Draghi JA, Plotkin JB, Andino R. 2015. Experimentally guided models reveal replication principles that shape the mutation distribution of RNA viruses. Elife 4:e03753.

[28] Boianelli A, Nguyen V, Ebensen T, Schulze K, Wilk E, Sharma N, Stegemann-Koniszewski S, Bruder D, Toapanta F, Guzmn C, Meyer-Hermann M. 2015. Modeling influenza virus infection: a roadmap for influenza research. Viruses 7(10):5274–304.

[29] Pinilla LT, Holder BP, Abed Y, Boivin G, Beauchemin CA. 2012. The H275Y neuraminidase mutation of the pandemic A/H1N1 influenza virus lengthens the eclipse phase and reduces viral output of infected cells, potentially compromising fitness in ferrets. Journal of Virology 86(19):10651–60.

[30] Holder BP, Beauchemin CA. 2011. Exploring the effect of biological delays in kinetic models of influenza within a host or cell culture. BMC Public Health 11(1):S10.

[31] Holder BP, Liao LE, Simon P, Boivin G, Beauchemin CA. 2011. Design considerations in building in silico equivalents of common experimental influenza virus assays. Autoimmunity 44(4):282–93.

[32] Paradis EG, Pinilla LT, Holder BP, Abed Y, Boivin G, Beauchemin CA. 2015. Impact of the H275Y and I223V mutations in the neuraminidase of the 2009 pandemic influenza virus in vitro and evaluating experimental reproducibility. PLOS One 10(5):e0126115.

[33] Banerjee S, Guedj J, Ribeiro RM, Moses M, Perelson AS. 2016. Estimating biologically relevant parameters under uncertainty for experimental within-host murine West Nile virus infection. Journal of the Royal Society Interface 13(117):20160130.

[34] Nguyen VK, Binder SC, Boianelli A, Meyer-Hermann M, Hernandez-Vargas EA. 2015. Ebola virus infection modeling and identifiability problems. Frontiers in Microbiology 6:257.

[35] Nguyen VK, Hernandez-Vargas EA. 2017. Windows of opportunity for Ebola virus infection treatment and vaccination. Scientific reports 7(1):8975.

[36] Malbec M, Porrot F, Rua R, Horwitz J, Klein F, Halper-Stromberg A, Scheid JF, Eden C, Mouquet H, Nussenzweig MC, Schwartz O. 2013. Broadly neutralizing antibodies that inhibit HIV-1 cell to cell transmission. Journal of Experimental Medicine 210(13):2813–21.

[37] Mateo M, Generous A, Sinn PL, Cattaneo R. 2015. Connections matter how viruses use cellcell adhesion components. Journal Cell Science 128(3):431–9.

[38] Xu Z, Waeckerlin R, Urbanowski MD, Van Marle G, Hobman TC. 2012. West Nile virus infection causes endocytosis of a specific subset of tight junction membrane proteins. PLOS One 7(5):e37886.

[39] Komarova NL, Anghelina D, Voznesensky I, Trinit B, Levy DN, Wodarz D. 2013. Relative contribution of free-virus and synaptic transmission to the spread of HIV-1 through target cell populations. Biology Letters 9(1):20121049.

[40] Iwami S, Takeuchi JS, Nakaoka S, Mammano F, Clavel F, Inaba H, Kobayashi T, Misawa N, Aihara K, Koyanagi Y, Sato K. 2015. Cell-to-cell infection by HIV contributes over half of virus infection. Elife 4:e08150.

[41] Dixit NM, Perelson AS. 2004. Multiplicity of human immunodeficiency virus infections in lymphoid tissue. Journal of Virology 78(16):8942–5.

[42] Sigal A, Kim JT, Balazs AB, Dekel E, Mayo A, Milo R, Baltimore D. 2011. Cell-to-cell spread of HIV permits ongoing replication despite antiretroviral therapy. Nature 477(7362):95.

[43] Komarova NL, Levy DN, Wodarz D. 2013. Synaptic transmission and the susceptibility of HIV infection to anti-viral drugs. Scientific Reports 3:2103.

[44] Lee CY, Kam YW, Fric J, Malleret B, Koh EG, Prakash C, Huang W, Lee WW, Lin C, Lin RT, Renia L. 2011. Chikungunya virus neutralization antigens and direct cell-to-cell transmission are revealed by human antibody-escape mutants. PLOS Pathogens 7(12):e1002390.

[45] Porta J, Prasad VM, Wang CI, Akahata W, Ng LF, Rossmann MG. 2016. Structural studies of chikungunya virus-like particles complexed with human antibodies: neutralization and cell-to-cell transmission. Journal of Virology 90(3):1169–77.

[46] Desmyter J, Melnick JL, Rawls WE. 1968. Defectiveness of interferon production and of rubella virus interference in a line of African green monkey kidney cells (Vero). Journal of Virology 2(10):955–61.

[47] Osada N, Kohara A, Yamaji T, Hirayama N, Kasai F, Sekizuka T, Kuroda M, Hanada K. 2014. The genome landscape of the African green monkey kidney-derived Vero cell line. DNA Research 21(6):673–83.

[48] Martins SD, Kuczera D, Ltvall J, Bordignon J, Alves LR. 2018. Characterization of dendritic cell-derived extracellular vesicles during dengue virus infection. Frontiers in Microbiology 9.

[49] Krejbich-Trotot P, Denizot M, Hoarau JJ, Jaffar-Bandjee MC, Das T, Gasque P. 2011. Chikungunya virus mobilizes the apoptotic machinery to invade host cell defenses. The FASEB Journal 25(1):314–25.

[50] Ramakrishnaiah V, Thumann C, Fofana I, Habersetzer F, Pan Q, de Ruiter PE, Willemsen R, Demmers JA, Raj VS, Jenster G, Kwekkeboom J. 2013. Exosome-mediated transmission of hepatitis C virus between human hepatoma Huh7. 5 cells. Proceedings of the National Academy of Sciences 110(32):13109–13.

[51] Zhang W, Jiang X, Bao J, Wang Y, Liu H, Tang L. 2018. Exosomes in pathogen infections: a bridge to deliver molecules and link functions. Frontiers in Immunology 12;9:90.

[52] Zhou W, Woodson M, Sherman MB, Neelakanta G, Sultana H. 2019. Exosomes mediate Zika virus transmission through SMPD3 neutral Sphingomyelinase in cortical neurons. Emerging Microbes & Infections 8(1):307–26.

[53] Boussier J. 2018. Chikungunya virus superinfection exclusion and defective viral genomes: Insights into alphavirus regulation of genetic diversity. PhD thesis. Universite Paris Diderot, Paris, France.

[54] Stapleford KA, Moratorio G, Henningsson R, Chen R, Matheus S, Enfissi A, Weissglas-Volkov D, Isakov O, Blanc H, Mounce BC, Dupont-Rouzeyrol M. 2016. Whole-genome sequencing analysis from the chikungunya virus Caribbean outbreak reveals novel evolutionary genomic elements. PLOS Neglected Tropical Diseases 10(1):e0004402.

[55] Schwarz MC, Sourisseau M, Espino MM, Gray ES, Chambers MT, Tortorella D, Evans MJ. 2016. Rescue of the 1947 Zika virus prototype strain with a cytomegalovirus promoter-driven cDNA clone. MSphere 1(5):e00246–16.

[56] Pastorino B, Bessaud M, Grandadam M, Murri S, Tolou HJ, Peyrefitte CN. 2005. Development of a TaqMan RT-PCR assay without RNA extraction step for the detection and quantification of African Chikungunya viruses. Journal of Virological Methods 124(1-2):65–71.

[57] Kirkwood TB, Bangham CR. 1994. Cycles, chaos, and evolution in virus cultures: a model of defective interfering particles. Proceedings of the National Academy of Sciences 91(18):8685–9.

[58] Nowak MA, Bonhoeffer S, Hill AM, Boehme R, Thomas HC, McDade H. 1996. Viral dynamics in hepatitis B virus infection. Proceedings of the National Academy of Sciences 93(9):4398–402.

[59] Efron B, Tibshirani R. 1986. Bootstrap methods for standard errors, confidence intervals, and other measures of statistical accuracy. Statistical Science 1:54–75.

